# Asymmetry in Histone Rotation in Forced Unwrapping and Force Quench Rewrapping in a Nucleosome

**DOI:** 10.1101/2020.10.21.348664

**Authors:** Govardhan Reddy, D. Thirumalai

## Abstract

Nucleosomes, the building blocks of chromosomes, are also transcription regulators. Single molecule pulling experiments have shown that nucleosomes unwrap in two major stages, releasing nearly equal length of DNA in each stage. The first stage, attributed to the rupture of the outer turn is reversible, occurs at low forces (≈ (3 - 5) pNs) whereas in the second stage the inner turn ruptures irreversibly at high forces (between ≈ (9 - 15) or higher) pNs. We show that Brownian dynamics simulations using the Self-Organized Polymer model of the nucleosome capture the experimental findings, thus permitting us to discern the molecular details of the structural changes not only in DNA but also in the Histone Protein Core (HPC). Upon unwrapping of the outer turn, which is independent of the pulling direction, there is a transition from 1.6 turns to 1.0 turn DNA wound around the HPC. In contrast, the rupture of the inner turn, leading to less than 0.5 turn DNA around the HPC, depends on the pulling direction, and is controlled by energetic and kinetic barriers. The latter arises because the mechanical force has to produce sufficient torque to rotate (in an almost directed manner) the HPC by 180°. In contrast, during the rewrapping process, HPC rotation is stochastic, with the quenched force *f_Q_* playing no role. Interestingly, if *f_Q_* = 0 the HPC rotation is not required for rewrapping because the DNA ends are unconstrained. The assembly of the outer wrap upon force quench, as assessed by the decrease in the end-to-end distance (*R_ee_*) of the DNA, nearly coincides with the increase in *R_ee_* as force is increased, confirming the reversible nature of the 1.6 turns to 1.0 turn transition. The asymmetry in HPC rotation during unwrapping and rewrapping accounts for the observed hysteresis in the stretch-release cycles in single molecule pulling experiments. Experiments that could validate the prediction that HPC rotation, which gives rise to the kinetic barrier in the unwrapping process, are proposed.

## Introduction

Genomes in eukaryotes, spanning roughly a meter long when stretched, are efficiently packed in a few micrometer sized cells. This spectacular process occurs routinely, resulting in the compaction of the genome accompanied by a density increase of over six orders of magnitude. The condensed material is chromatin, - a polymer whose building block is the nucleosome,^1^ made up of a complex between DNA and the highly conserved histone proteins. The ≈ 147 base pairs (bps), roughly the persistence length of DNA, are wrapped around the (≈ 10 nm) octameric histone protein core (HPC) (Figure 1). It follows that DNA is tightly wound around the histones ≈ (1.6 − 1.7) times (Figure 1). Key cellular processes, such as replication, recombination and transcription depend in part on the positioning and the dynamics of the nucleosomes.^2,3^ Because the primary function of nucleosomes is the regulation of transcription,^4^ it is important to understand the forces that govern the stability of the nucleosome, and the mechanism by which DNA unwraps and rewraps.

**Figure 1:**
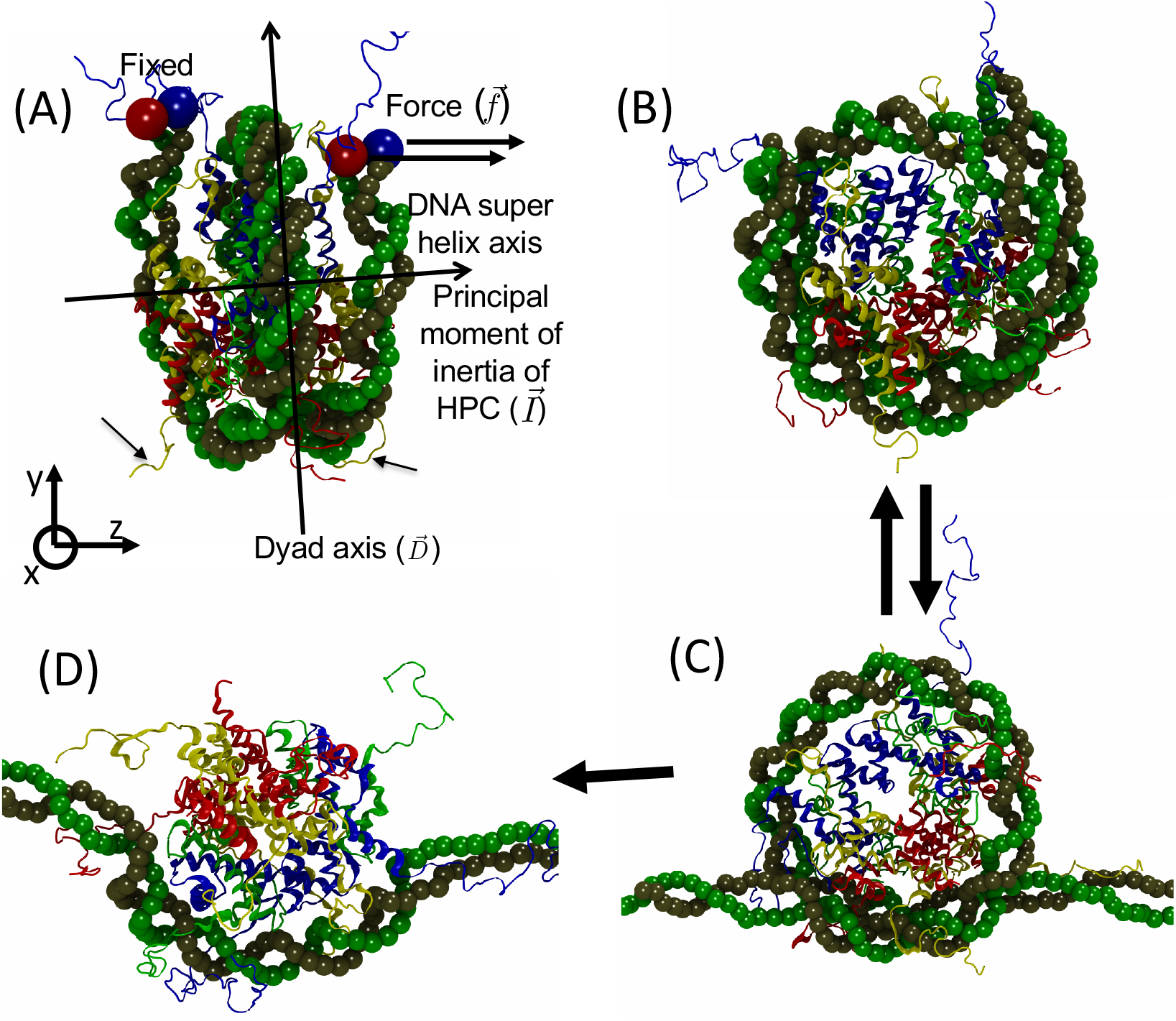
A nucleosome has a double stranded DNA, which wraps around the HPC. DNA strands are shown in tan and green color. Two copies each of the four proteins H2A, H2B, H3 and H4 form the HPC. H2A, H2B, H3 and H4 are colored in yellow, red, blue and green, respectively. (A) Nucleosome orientation at the start of the simulations. The two fold nucleosome dyad axis, 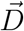, passing through the center of the wrapped DNA, is along the *y* direction and lies in the *xy* plane. The *xy* plane, through the center of the figure and perpendicular to the plane of the paper roughly separates one copy of the H2A, H2B, H3 and H4 protein complex from the second. The DNA super helix axis and the principal moment of inertia 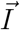 of the HPC point along the *z* direction and are in the *xz* plane. The two ends of the double stranded DNA are shown as red and blue beads. One end of the DNA is fixed and a constant force is applied to the other end, either along the *x* or the *z* direction. The small slanted arrows point to the histone protein tails, which interact with the DNA. When force is applied to the ends of the DNA, the first reversible structural transition occurs shown as (B) to (C). The outer DNA turn unwraps from the HPC. The second abrupt irreversible structural transition occurs between (C) and (D), where the inner turn unwraps. The figure is rendered using Visual Molecular Dynamics (VMD).^17^

Single molecule pulling experiments on chromatin fibers,^5–11^ and arrays of nucleosomes and a single nucleosome^5,12–16^ have given quantitative insights into their nanomechanics. Of particular relevance here are the pulling experiments on single nucleosomes using the Widom 601 sequence,^12^ which established that the fully wrapped DNA unravels in two major stages upon application of mechanical force, *f* (Figure 1). In the first stage, occurring at low forces, *f* ≈ (3 − 5) pN, the outer turn of the DNA “rips” or unwraps releasing ≈ 21 ± 0.22 nm (roughly 60 - 70 base pairs) length of DNA.^12–14^ In the force range (f ≤ 5 pN), DNA hops between the fully wrapped and partially unwrapped states. Thus, it is surmised that the observed two-state hopping is an equilibrium reversible process. ^12,14^ The low force transition is interpreted as peeling of the outer turn in which approximately 1.6 to 1.0 turn of DNA is unwound (Figure 1B to 1C). The second unwrapping stage, which occurs in the force range *f* ≈ (9 − 15) pN or more ≈ (15 − 20) pN, is attributed to an irreversible peeling of the inner turn of the DNA.^12,14^ During this transition there is an additional release of ≈ 22 ± 0.18 nm of DNA, which results in unwinding from 1.0 to less than 0.5 turn (Figure 1C to 1D). The DNA release during this stage is abrupt, ^5^ and shows signs of hysteresis, inferred from the finding that the force extension curves (FECs) during the stretching/release of force do not coincide.^12^ We refer to the three dominant states of the wrapped DNA as 1.6**N**, 1.0**N**, and 0.5**N**, where **N** stands for nucleosome and the number in front is the amount of DNA turns around the HPC. The numerical values specifying the number of turns are approximate.

High forces required to peel the inner wrap is larger than estimates using equilibrium arguments.^12,18,19^ To explain this behavior, it was conjectured that there could be electrostatic attraction between DNA (negatively charged) and the positively charged HPC.^5,12^ However, neither the crystal structures^20,21^ nor the equilibrium accessibility of the nucleosomal DNA provides evidence for such a patch that may be conducive to large favorable interactions between the histones and the partially wrapped DNA. ^22^ This observation prompted Kulic and Schiessel (KS),^22^ in a remarkable paper, to propose an alternative explanation. DNA is negatively charged and when the DNA is fully wrapped around the nucleosome, there could be considerable electrostatic repulsion between the inner and outer wraps of the DNA. This unfavorable interaction results in a small free energy barrier observed during the first stage of DNA unraveling. After the first stage of unraveling, the nucleosome transitions to the 1.0**N** state. Consequently, the unfavorable DNA-DNA electrostatic repulsion is muted, and the structure of the nucleosome is stabilized, which results in a much higher free energy barrier (and hence larger values of f) for unraveling the inner wrap of the DNA. These generic arguments suggest that the global unwrapping mechanism may not be sensitive to the DNA sequence. We hasten to point out that not all nucleosomes are physically identical. The post translational modifications chemically alter the histones. In addition, DNA nanomechanics and dynamics are sequence dependent.^23–25^

Given the importance of HPC in the genome organization, it is not surprising that there are many theoretical and computational studies^18,19,22,26–34^ probing the stability of nucleosomes. In some of these studies the DNA is modeled as a semi-flexible polymer, and the HPC as a rigid spherical bead or a helical spool. ^22,30^ Such a description is remarkably successful in predicting the two major stages of DNA unwrapping. However, they are not entirely successful in fully explaining the equilibrium nature of the first stage of unwrapping and the kinetically controlled (manifested as hysteresis in the rewrapping of DNA from the 0.5**N** state upon force quench) second transition. Some of the reasons could be that the theoretical studies assume thermodynamic equilibrium during nucleosome DNA pulling, where as the energy barriers could exist for kinetic reasons. In addition to the theories based on DNA elasticity, coarse-grained simulations have been used to explore the free energy landscape of nucleosomes. ^32,33^ In a study that is related to our investigation^33^ force-dependent free energy profile of the DNA as a function of the extension was used to provide quantitative estimates of the unwrapping forces.

Most, if not all, of the computational studies are DNA centric with scant attention to the role that HPC plays in the unwrapping or rewrapping processes. In order to remedy this situation, we investigate forced-unwrapping and force-quench rewrapping of a single nucleosome using a coarse grained (CG) model of the nucleosome and Brownian dynamics simulations. In accord with experiments, we find that there is a reversible transition from 1.6**N** to 1.0**N** states followed by a transition to the 0.5**N** state at high forces. Upon force quench, we find that there is hysteresis in the 0.5**N** → 1.6**N** transition. We show that hysteresis observed in the pulling experiments is due to the distinctly asymmetric role the external mechanical force plays in the DNA unwrapping and rewrapping pathways. In the second irreversible transition (1.0**N** → 0.5**N**), kinetic barriers arise because sufficient torque has to be generated by the mechanical force to rotate the HPC by nearly 180°, which is resisted by the histone tails. In contrast, rewrapping of the DNA onto the HPC during the second transition upon quenching the force to low values (≤ 3pN), occurs only when the principal moment of inertia of the HPC aligns along the pulling direction. The alignment occurs stochastically by thermal fluctuations. The external force does not play a role in aligning the HPC and the DNA. This asymmetry in the role played by HPC rotation dramatically alters the rewrapping pathways, thus contributing to the observed hysteresis in single molecule pulling experiments. We show that the requirement for HPC rotation to unwrap the DNA increases the barrier has both an energetic and a kinematic effect. The latter renders the rupture of the inner turn irreversible. We predict that the time scale to unwrap the inner turn should increase exponentially with the solvent viscosity, which is amenable to experimental test.

## Methods

### SOP model for the nucleosome

The simulations use the Self-Organized Polymer (SOP) model, ^35^ which has proved to be accurate in describing the dynamics of polymerase-induced transcription bubble formation,^36^ folding of large proteins,^37^ effect of force on RNA,^35^ and cation-dependent folding of a number of RNA molecules.^38^ Each residue in the histone is represented using a single bead. Following our earlier work, ^36^ we represent the nucleotides in the DNA using a single bead, which is sufficient to characterize the mechanical properties (for example persistence length (*l_p_*) and salt-dependent changes in *l_p_*) of DNA accurately. Predictions using the model^36^ for bubble formation in promoter DNA during bacterial transcription have found experimental support, ^39,40^ thus validating the model.

In the SOP model, the center of a bead representing the protein residue is at the *C_α_* atom, and the bead representing the nucleotide is at its center of mass. The SOP model for the nucleosome is constructed using the crystal structure^20^ of the human nucleosome available in the Protein Data Bank (PDB ID: 2CV5). The 147 base pairs palindromic DNA sequence from the human *α*-satellite DNA^21^ binds symmetrically to the HPC. The DNA is split into two halves containing 73 and 72 bps, with a single base pair located on the dyad. We performed Brownian dynamics simulations^41^ at temperature, *T* = 300 K to determine the mechanism of force-induced unwrapping and force-quench rewrapping of DNA from the nucleosome. Details of the model and the simulation methodology are given in the Supplementary Information (SI).

## Results

Most of the results are obtained without the histone tails, whose role is examined to ensure that the overall unwrapping mechanism is unaltered. In the KS study^22^ and subsequent generalizations, it was shown that many (but not all) aspects of tension-induced unwrapping of the nucleosome could be explained using a purely elastic model of the wrapped DNA. To go beyond the elastic model description of DNA and rigid body cylinder representation of the HPC, we use the SOP model for the DNA-protein complex to elucidate the molecular details of dynamics of unbinding of DNA from the nucleosome with focus on the HPC dynamics, which affect the DNA rupture and rewrapping upon force quench.

### DNA unwraps in two stages

The orientation of the nucleosome at the start of the simulation is shown in Figure 1A. The principal moment of inertia of the HPC (see SI), 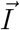, is oriented along the *z* direction. In this orientation, the end-to-end vector of the DNA, 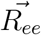, points along the *x* direction (Figure 1A). One end of the double stranded DNA is fixed, and a constant force, 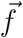, is applied to the other end on the 2 beads of the double stranded DNA. When the applied 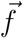 is parallel to 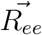 (along the *x* direction), it is readily transmitted to the rest of the DNA.

In accord with the experiments and previous simulations using a different coarse-grained model,^33^ we find that DNA unwraps in two stages^12^ (Figure 2). It can be concluded that the reversible unwrapping of the outer turn followed by a non-equilibrium rupture of the inner turn, representing the global two stage rupture of DNA, are likely to be independent of the DNA sequence because two different sequences yield the same pattern of DNA rupture. In all the trajectories, the 1.0**N** state is observed as long as the force is applied along the *x* direction (parallel to 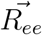), implying that it is an essential intermediate in the forced-rupture of the nucleosome. Pulling experiments^12^ find that a transition from 1.6**N** turns to 1.0**N** turn occurs in the first stage (Figure 2A(i) to 2A(ii)/(iii)). Our simulations show that the peeling of the outer wrap occurs at a force of ≈ 5 pN (see the time dependent changes in *R_ee_*(*t*) in Figure 2). During this transition the extension of the DNA, *R_ee_*, increases by ≈ (21 − 22) nm, with the experimental value being about 21 nm. The second rip, which occurs when *f* exceeds ≈ 10 pN, involves a transition to the 0.5**N** state from the 1.0**N** state (Figure 2A(ii) to 2A(iv)). The increase in extension in the second transition is ≈ 25 nm, which is also in agreement with 22 nm measured in experiments. In our simulations, only when *f* exceeds 25 pN there is an irreversible 1.0**N** → 0.5**N** transition on the time scale of simulations (tens of *μ*secs) (Figure 2A(ii) to 2A(iv)).

**Figure 2:**
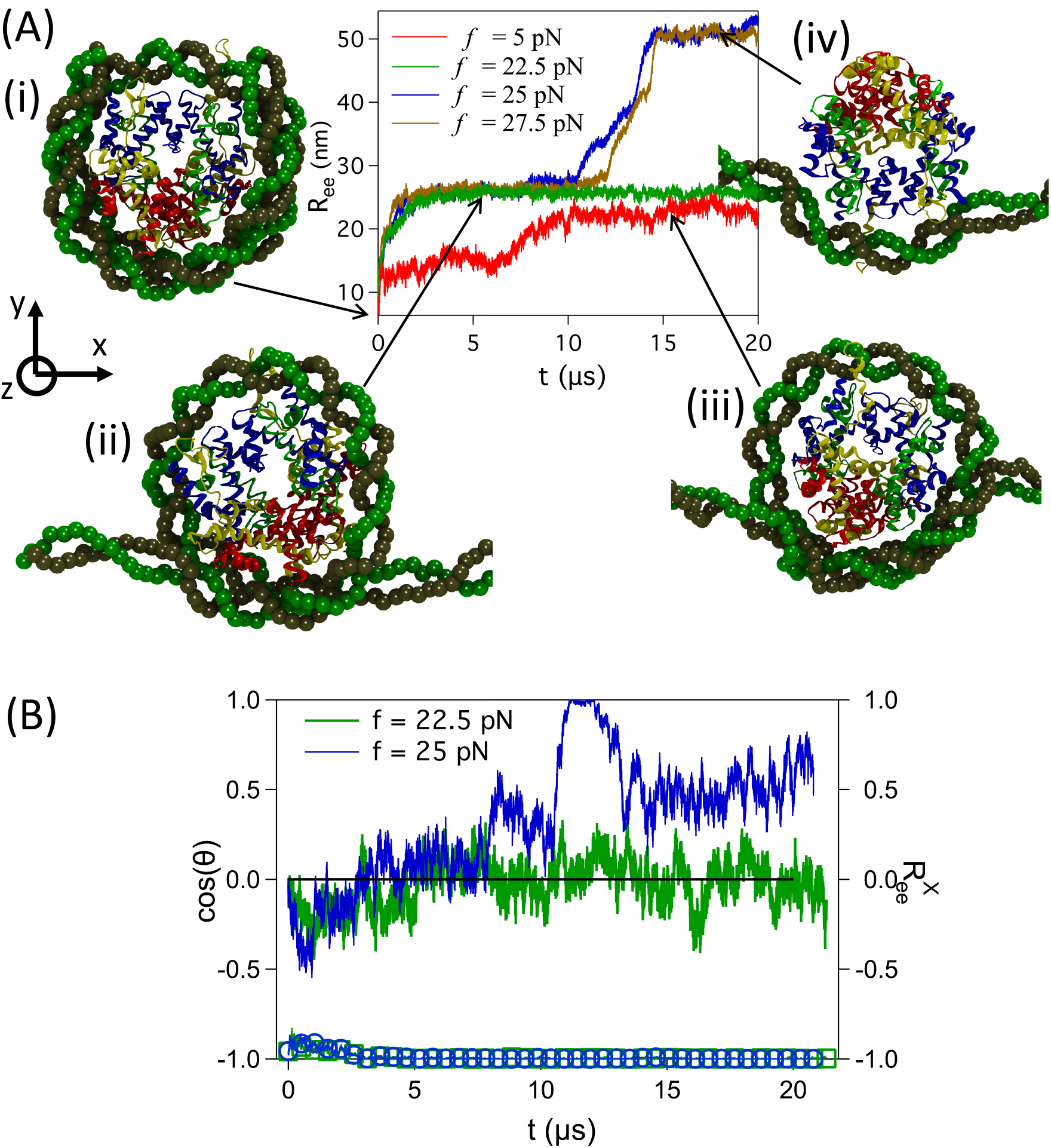
(A) Stages in DNA unwrapping when the force 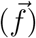 is applied along the *x* direction. In the initial conformation, the dyad axis of the nucleosome 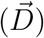, principal moment of inertia of the HPC 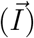, and DNA end-to-end distance 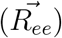 point along the *y*, *z* and *x* directions, respectively. *R_ee_* is plotted as a function of time (*t*) at different values of the constant *f*. The sequence of structural changes of the nucleosome that occur in two stages are shown in (i) to (iv). (B) The 1.0**N** state is observed at the end of the transition for *f* = 5 pN (red curve in (A)) at *t* ≈ 20 *μs* and at *t* ≈ 5 *μs* for *f* = 22.5 pN (green curve in (A)). The value of cos(*θ*) fluctuates around 0 (green), since *θ* ≈ 90°. At *f* = 25 pN, DNA unwraps completely from the HPC resulting in the transition from 1.0**N** to 0.5**N** state (blue curve in (A)), which requires a 180° rotation by the HPC. This rotation is observed between 10 − 15 *μ*sec (see the jump in cos(*θ*)). For both *f* = 22.5 pN (blue circles) and 25 pN (green squares), the x-component of the unit of vector of *R_ee_*, 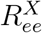 is ≈ −1, indicating that *R_ee_* is along the *x* direction and does not change orientation during the transitions.

The findings in the simulations are in reasonable accord with experiments. In the pulling experiments, ^12^ the first transition is centered at ≈ 3 pN and whereas the second rupture has a most probable value of ≈ 9 pN. However, the distribution of measured rupture forces is broad. In our simulations, the corresponding forces are ≈ 5 pN and ≈ 25 pN, respectively. It is worth noting that forces in the range of ≈ (15 − 30) pN might be required to unwrap the inner turn^12,14^ depending on the loading rate. Considering that we did not choose the model parameters to reproduce any feature of the pulling experiments, we surmise that the SOP simulations semi-quantitatively reproduce the two stage DNNA unwrapping found in experiments.

The larger average rupture forces, compared to experiments, for the second transition is due to the limitations on the length of the simulations. The traces are on a time scale of seconds in the experiments, whereas the maximum time scale achievable, even using the coarse-grained SOP model, is on only the order of tens of microseconds. Therefore, it is necessary to use high forces (by implication high pulling speeds) in order to observe the unwrapping events in about (10 − 20) *μs*. The most probable unbinding force for the second non-equilibrium transition (see below) is expected to scale as ln(*v*),^42^ where *v* is the velocity with which one end of the protein is pulled keeping the other end fixed. Therefore, we believe that the unwrapping mechanism would not change significantly even if the mean rupture force for the second transition in the simulations is larger than in the experiments.

### Forced unwrapping induces HPC rotation

Pulling experiments and computational studies have largely focused on the forced-unwrapping of the DNA. How the HPC responds to the forces applied to DNA is unknown. It is clear from the KS model, ^22^ and the early studies by Cui and Bustamante^6^ that HPC must rotate during the rupture of the inner turn, although the details of how this transpires cannot be elucidated using the cylinder representation of the HPC or from experiments. We find that during the 1.0**N** → 0.5**N** transition, the HPC undergoes a 180° rotation in agreement with the previous studies. ^6,22^ As the HPC rotates, the direction of the principal moment of inertial vector associated with the HPC, 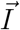, which initially points in the *z* direction, undergoes 180° rotation in the *xz*-plane around the *y* direction. In contrast, 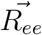, which was along the *x* direction does not undergo any change in orientation.

The HPC rotation is quantified using cos(*θ*), where *θ* is the angle between the vectors 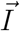 and 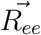 (see Eqs. (S3) and (S4) in the SI for details). The angle *θ* varies from either −90° to 90° or 90° to −90° during the 180° rotation of the HPC in the *xz*-plane (Figure 2B). The abrupt jump in cos(*θ*) (from ≈ 0 to ≈ +1 in Figure 2B) is evidence of HPC rotation during the unwrapping of the inner turn. In contrast, the *x*-component of the end-to-end unit vector, 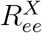, is pinned at ≈ – 1 during the HPC rotation indicating that the direction of 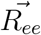 does not change during the 1.0**N** → 0.5**N** transition (Figure 2B). The torque required for the rotation of the HPC increases the (kinetic) barrier for the unwrapping of the inner turn (see below for a rough estimate).

### Outer turn unwrapping transition is reversible

In order to demonstrate that the peeling of the inner wrap of the DNA is reversible, we applied 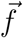 along the *z* direction (perpendicular to 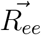) on one end of the DNA while keeping the other end fixed. The stretching energy is 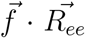 with 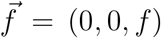. The nucleosome was aligned in the same orientation as before (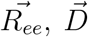 and 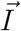 approximately point along the *x*, *y*, and *z* directions, respectively) (Figure 3A(i)). In this geometry, 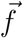, which is perpendicular to 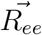 does not propagate along the contour of the DNA. As before the 1.6**N** → 1.0**N** transition occurs at *f* ≈ 5 pN. The time dependent changes in *R_ee_*(*t*) in both the pulling geometries are almost identical (compare the red curves in Figures 2 and 3). The independence of the unwrapping of the outer turn on the pulling geometry suggests that this transition is reversible.

**Figure 3:**
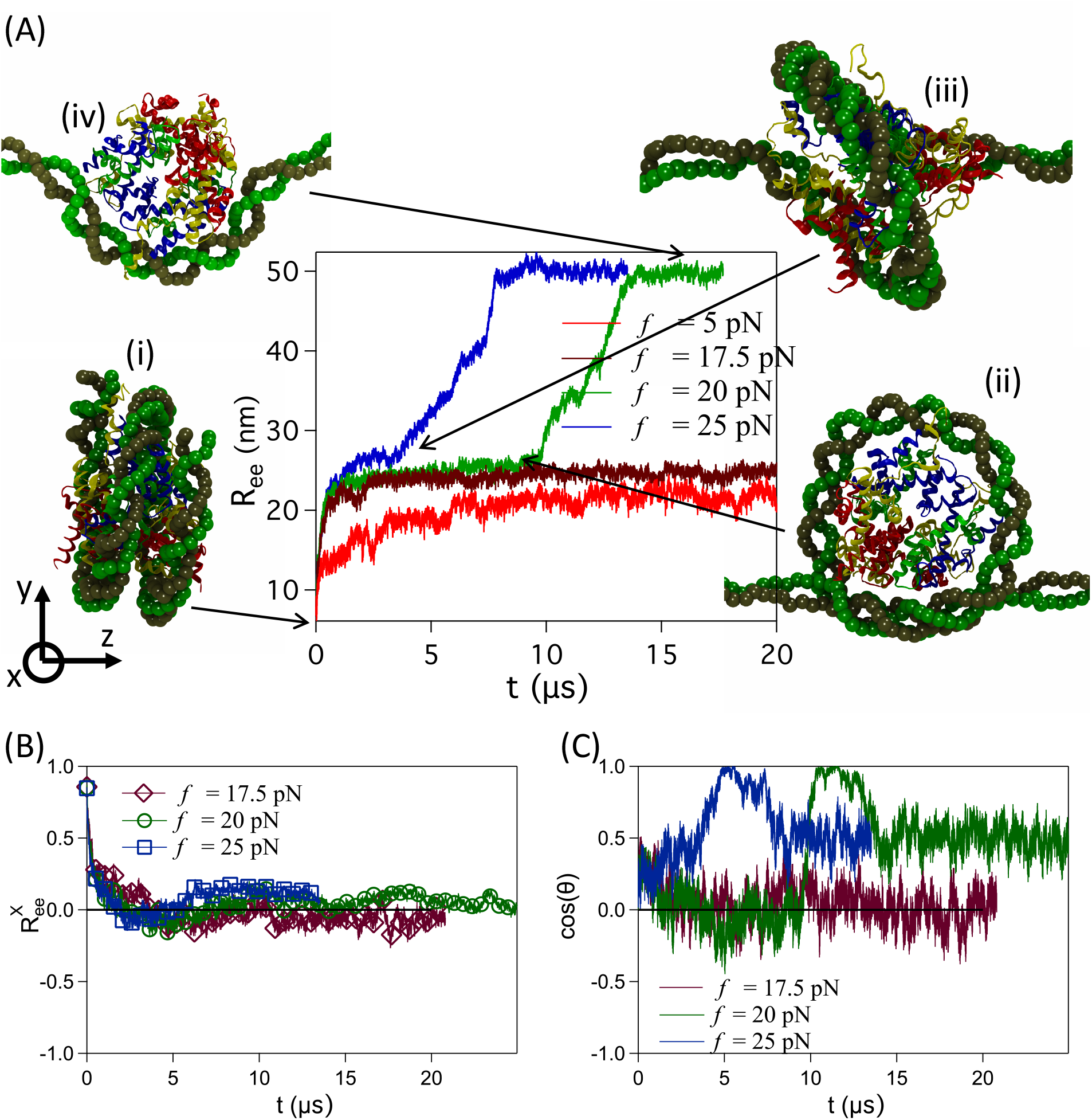
Pulling DNA along the *z*-direction. (A) At *t* = 0, 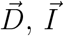 and 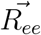 are along the *y*, *z*, and *x* directions, respectively. *R_ee_* is plotted as a function of *t*. (B) In the initial 2 *μsec*, as 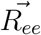 aligns along 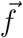, the direction 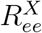 decays to ≈ 0. (C) At *f* = 17.5 pN, 1.0**N** state is observed at the end of the first transition. On this time scale cos(*θ*) fluctuates around 0. At *f* = 20 pN, DNA unwraps completely from the HPC. The transition from 1.0**N** to 0.5**N** state, which requires a 180° rotation by the HPC occurs between 10 − 15 *μ*sec (cos(*θ*) jumps abruptly in the green trajectory). At *f* = 25 pN, DNA continuously unwraps from the HPC as it rotates with the jump in cos(*θ*) occurring on shorter times.

The 1.0**N** state is observed for *f* in the range 5 pN ≤ *f* ≤ 20 pN (Figure 4). During the 1.6**N** → 1.0**N** transition, the whole nucleosome rotates by 90° in the *xz* plane about the *y* direction as the applied force realigns 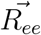 along the force direction (*z* direction) (Figure 3A(i) to 3A(ii) and 3B). Upon rotation, the vectors 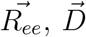 and 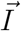 of the nucleosome point along the *z*, *y*, and *x* directions, respectively (Figure 3A(ii)). As a consequence, 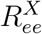 decreases from ≈ 1 to ≈ 0 in ≈ 2.0 *μsec* after the application of force (Figure 3B). We do not detect the 90° rotation of the nucleosome from the plots of cos(*θ*) as both 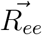 and 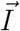 simultaneously undergo 90° rotation around the *y* direction (Figure 3C).

**Figure 4:**
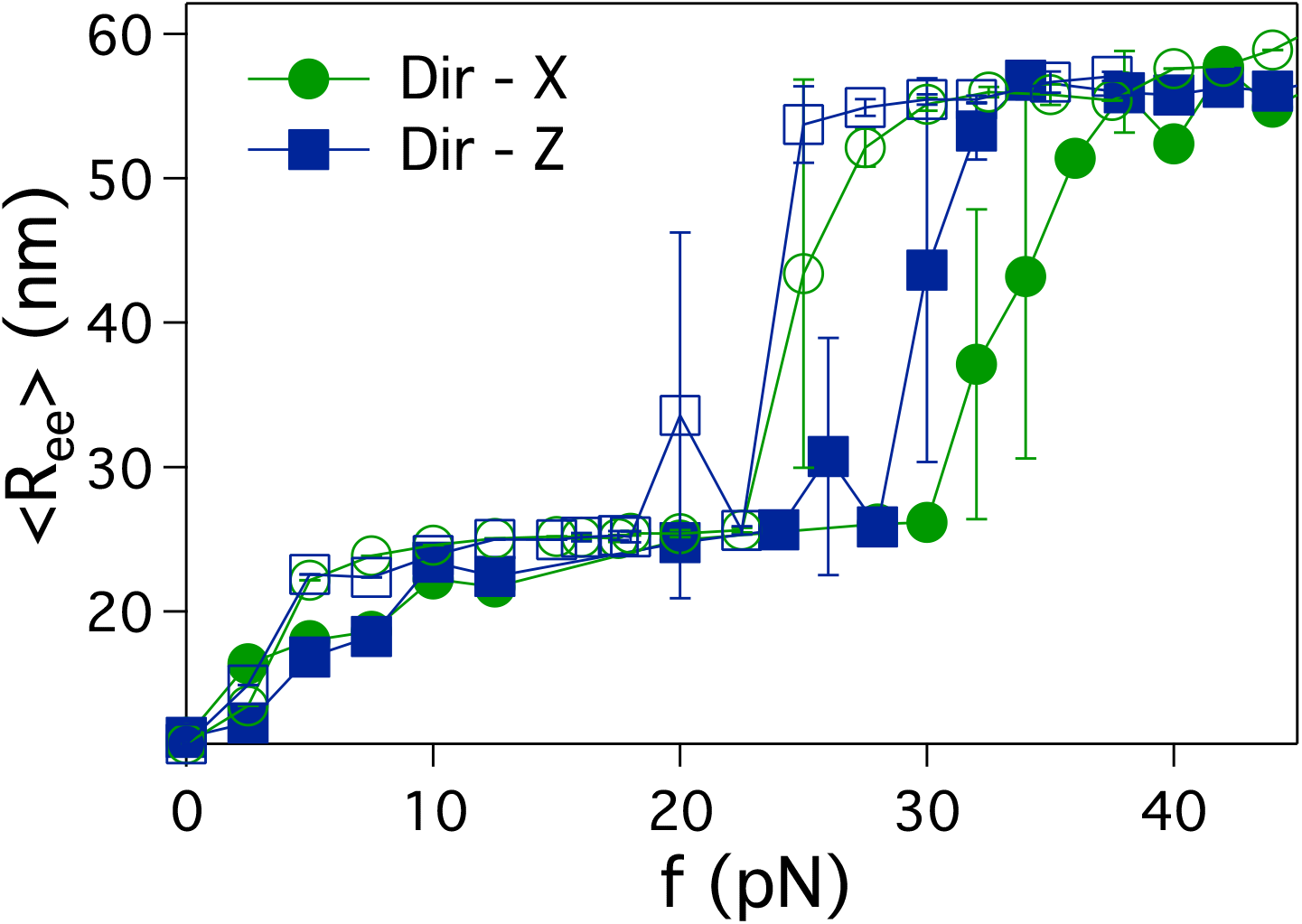
Average *R_ee_*, 〈*R_ee_*〉 plotted as a function of *f*. The open (closed) symbols represent results without (with) histone tails. The superposition of all the curves at low forces once again shows that the unwrapping of the outer turn is reversible. The charged residues in the histone tails interact with the DNA and prevent the rotation of the HPC. DNA unwraps from the HPC at higher forces compared to when the histone tails are absent. Inclusion of the histone tails does not alter the global unwrapping mechanism. For each data point the DNA is pulled at constant *f* for at least 6 ×10^8^ Brownian dynamics simulation time steps, which is 15 *μ*s.

### Population of the 1.0N state depends on both the magnitude of force and the pulling direction

We next performed simulations at *f* = 20 pN applied along the *z* direction. The time dependence of *R_ee_*(*t*) (the green line in figure 3A) shows the familiar two stage unwrapping transition. The lifetime of the 1.0**N** state is at least ≈ 10 *μs* (Figure 3). Such a long lifetime suggests that for the 1.0**N** → 0.5**N** transition there must be a free energy barrier because the HPC must rotate by 180°, as shown earlier (see Figure2B) when the force was applied along the *x* direction (Figure 3C). During the HPC rotation, only the moment of inertia 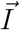 rotates by 180°, while 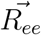, which points along the *z* direction does not change orientation even during the inner unwrapping transition.

In contrast, when *f* = 25 pN is applied perpendicular to 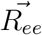, the 1.0**N** state is not detected in the transition to the 0.5**N** state (see the blue line in Figure 3). Comparison of the structures, Figure 3A(ii) and Figure 3A(iii), shows dramatic differences. At this force, 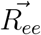 become parallel to 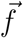 in the initial ≈ 2 *μsec* (Figure 3B). In this short time, more than 1.0 turn of DNA is unwrapped. As a consequence, the 1.0**N** state is not populated in this pathway (Figure 3A(iii)). We find that at very large forces there is a continuous transition from the 1.6**N** to 0.5**N** state bypassing the 1.0**N** state (Figure 3A(iii),3B,3C).

### Role of histone tails

To ensure that the two-stage unwrapping mechanism is robust, we performed simulations to assess the role of histone tails (see SI for details). We find that the unwrapping mechanisms are similar with and without histone tails. However, due to additional favorable interactions between the histone tails and the DNA, the magnitudes of the rupture forces, especially for the second transition (1.0**N** → 0.5**N**), are higher. The average extensions in *R_ee_* at the end of the transitions, as a function of the applied *f* along the *x* direction (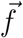 parallel to 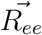) and *z* direction (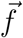 perpendicular to 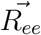) are shown in Figure 4. An external force between (5 − 8) pN is required to observe the first transition irrespective of the direction of the applied force. The dependencies of *R_ee_* on *f* are similar at forces at which outer unwrapping is observed, resulting in the population of the 1.0**N** state.

In the presence of histone tails, a higher magnitude of force is required to observe the 1.0**N** → 0.5**N** transition because they are present on either side of the DNA (see Figure 1A), leading to favorable electrostatic interactions with the DNA. As a result, the 180° rotation of the HPC, required for the second transition, is further hindered. Apart from the differences in the rupture forces the nature of the two stage transition is not qualitatively affected.

### Differences in wrapping and unwrapping mechanisms explain hysteresis

In order to determine the differences in the assembly (rewrapping from the stretched 0.5**N** state to the 1.6**N** state) and forced-unwrapping of DNA, we performed force quench simulations. The nucleosome is first prepared in the 0.5**N** state by applying a constant force, and subsequently the force is quenched to *f_Q_* = 0 (may be difficult to realize in experiments). Note that the direction of pulling does not affect the formation of the fully unwrapped (0.5 **N**) state. If the transitions were reversible, we would expect the assembly pathway 0.5**N** ↔ 1.0 **N** ↔ 1.6**N**. Instead, we find that rewrapping mechanism upon force quench to *f_Q_* = 0 pN, (Figure 5) is dramatically different from the force-induced unwrapping pathway (Figure 2). When *f_Q_* = 0, the HPC rotation does not occur because in the DNA ends are unrestricted in the simulations. As a result, DNA undergoes stochastic fluctuations as it rewraps around the HPC (Figure S2). The two free ends of the DNA slide past one another in order to wrap around the nucleosome (Figure 5(iii) and 5(iv)). However, this wrapping pathway is prohibited in single molecule pulling experiments as the ends of the DNA are pinned by attaching them through a handle to a polystyrene bead held in an optical trap.^12^ Hence, the DNA ends cannot readily wrap around the HPC.

**Figure 5:**
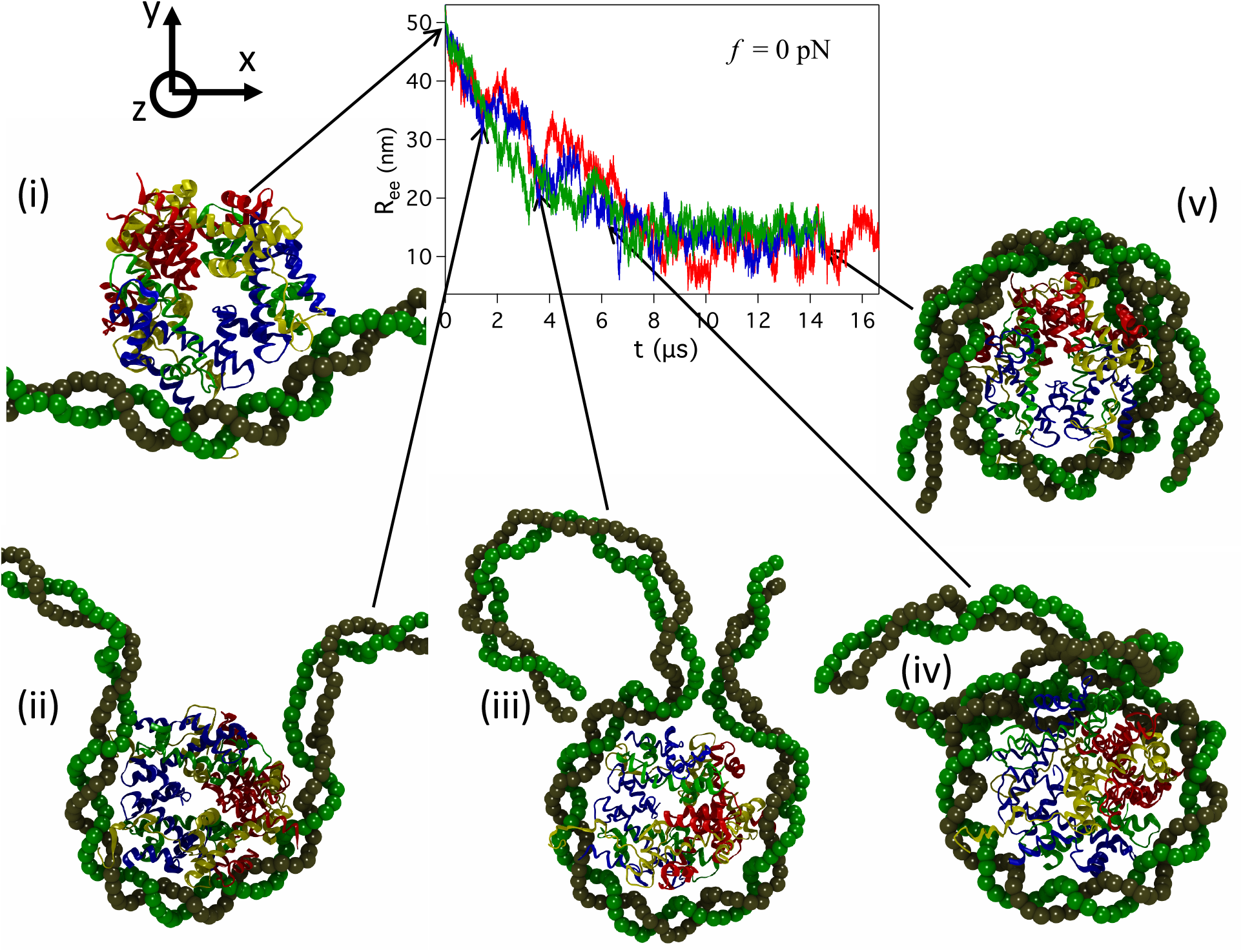
Rewrapping of DNA around the HPC upon force quench. Displayed are the structural transitions at select time points, in the pathway to form a fully wrapped nucleosome after quenching the force to *f_Q_* = 0 pN. The time-dependent decrease in *R_ee_* in the three different trajectories, shown in red, blue and green, are superimposable.

The situation in the optical trap experiments can be mimicked in the simulations if *f_Q_* ≠ 0. In such a setup, DNA can rewrap around the HPC only when 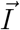 of the HPC aligns with the direction of 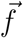 (*x* direction) (see Figure 6(iii)). The time dependent decrease in *R_ee_*(*t*) in the representative rewrapping trajectory for *f_Q_* = 3 pN shows that upon force quench, *R_ee_*(*t*) decreases by ≈15 nm (the green curve in Figure 6) rapidly. The structures that are sampled during the long time period when there is a plateau in both *R_ee_* (*t*) and cos(*θ*) correspond to the 0.5**N** state. The time dependent changes in *R_ee_*(*t*) further shows that 0.5**N** → 1.0**N** transition, which takes place between 30 - 40 *μs* is the key event in the assembly process. It should be pointed out that at *t* ≈ (30 – 40) *μs* the value of *R_ee_* is similar (the green curve in Figure 6) to that found after outer turn forced unwrapping (see Figure 2), implying that the 1.0**N** state is sampled during unwrapping and rewrapping processes. The formation of the 1.0**N** state both during unwrapping and upon force quench affirms that the first transition is reversible. The 1.0**N** → 1.6**N** transition leading to the complete rewrapping of the DNA on the HPC occurs rapidly when the jump in cos(*θ*) is complete.

**Figure 6:**
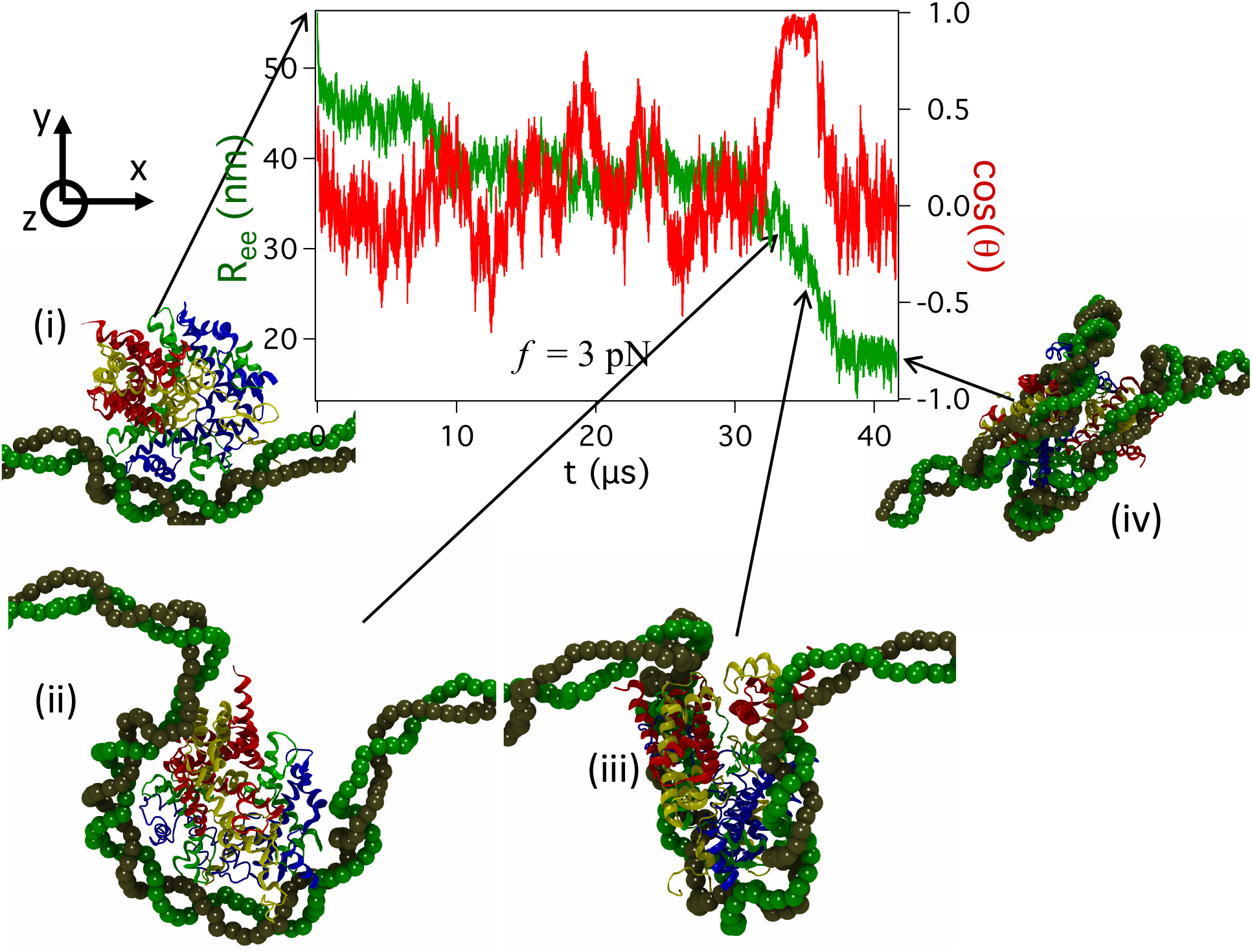
Rewrapping dynamics. *R_ee_* and cos(*θ*) are plotted as a function of *t* for a DNA folding trajectory for *f_Q_* = 3 pN.

Note that prior to the 1.0**N** → 1.6**N** transition, cos(*θ*) jumps to unity (red curve in Figure 6) before decreasing to zero when the nucleosome with the fully wrapped DNA reforms. Naively, it may appear that the spike in cos(*θ*) is the reverse of event that occurs when unwrapped at high forces (see the blue curve in Figure 2B). However, in the latter case the sudden increase in cos(*θ*) is directed in the sense that the mechanical force generates a torque along a specific direction causing the observed rotation. In sharp contrast, in the reassembly process the alignment of 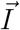 of the HPC along the direction of 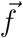 occurs stochastically due to thermal fluctuations of the HPC (Figure 6(iii)). Thus, in the process of rewrapping of the DNA on HPC, force does not play any role in aligning 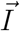 of the HPC along the 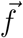 direction. The applied 3 pN force does not generate torque in the direction opposite to what was observed during DNA unwrapping by force. Alignment of HPC in a favorable orientation for DNA wrapping is a random event. The different role *f* plays in the wrapping and unwrapping pathways of the DNA contributes to the observed hysteresis in the force-extension curve for the nucleosome. ^12^ Thus, inherently the kinetically driven second unwrapping transition exhibits non-equilibrium signatures. Similar DNA wrapping mechanism is also observed for force quenches to *f_Q_* = 2 and 4 pN (Figure S3).

### Rotational barrier in the unwrapping of the inner turn

In order to fully unwrap the DNA from the populated 1.0**N** state, the magnitude of *f* must be sufficiently large to generate the needed *torque to rotate* the HPC by 180° in water. In order to account for histone rotation, we modified the KS^22^ theory, which models DNA as a wormlike chain (WLC) of length, *L*, and stiffness *κ*. The HPC is modeled as a cylinder with radius, *R*. The WLC is adsorbed on the cylinder in a helical manner with a pitch, *H*. The torsional stiffness of the WLC is neglected because the ends of the DNA are free to rotate. As force *f* is applied to the WLC (Figure S1), the WLC bends and desorbs from the cylinder and simultaneously causes the cylinder to rotate. The desorption angle *α* describes the length of the WLC adsorbed on the cylinder (*α* = 0 and *π* correspond to the 1.0**N** state, and nearly completely unwrapped state, respectively). The angle *β*, accounting for the degree of rotation of the cylinder, describes the orientation between the axis of the cylinder, 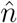, and the *z* direction (Figure S1). Due to the symmetry in the system, the axis of the cylinder, 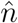, rotates only in the *yz* plane. The total energy of the mechanical representation of the nucleosome is,

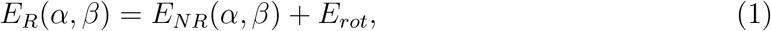

where *E_NR_*(*α*, *β*)^22^ is reproduced in the SI.

### Estimation of *E_rot_*

We calculated the contribution of the barrier to HPC rotation, *E_rot_*, using the elastic model. The energy required to rotate the cylinder through an angle *β* by the torque, *τ*(*α*, *β*′), generated due to *f* acting on the WLC arms is,

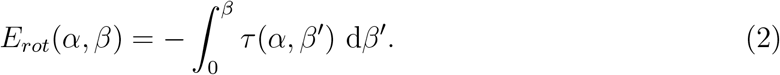

The generated torque increases the transition energy barrier for the nucleosome to go from the 1.0**N** state (*α* = 0) to the state where DNA is completely unwrapped (*α* = *π*).

Plots of *τ*(*α*, *β*) (Eq. S13) and *E_rot_*(*α*, *β*) (Eq. S14), calculated numerically as a function of *α*, subject to the constraint (Eq. S8) for different *f* are shown in Figure S4 (details in the SI). The addition of *E_rot_*(*α*, *β*) to the energy of the WLC and cylinder system increases the transition barrier to fully unwrap the DNA (Figure 7A). For *f* = 0.8 *k*_B_*T*/nm, the barrier for WLC unwrapping from the cylinder increases by ≈ 2.5 *k*_B_*T* due to *E_rot_*(*α*, *β*) (Figure 7A). The energy difference between the DNA wrapped state and unwrapped state, Δ*E_R_*(= *E_R_*(*α* ≈ *π*) – *E_R_*(*α* ≈ 0)) and Δ*E_NR_*(= *E_NR_*(*α* ≈ *π*) – *E_NR_*(*α* ≈ 0)), as a function of *f* shows that the requirement to rotate the cylinder increases the transition barrier, and the energy difference between the states *α* ≈ *π* and *α* ≈ 0 for all *f*. Thus, a larger *f* than previously estimated would be needed to unwrap the DNA from the cylinder (Figure 7B).

**Figure 7:**
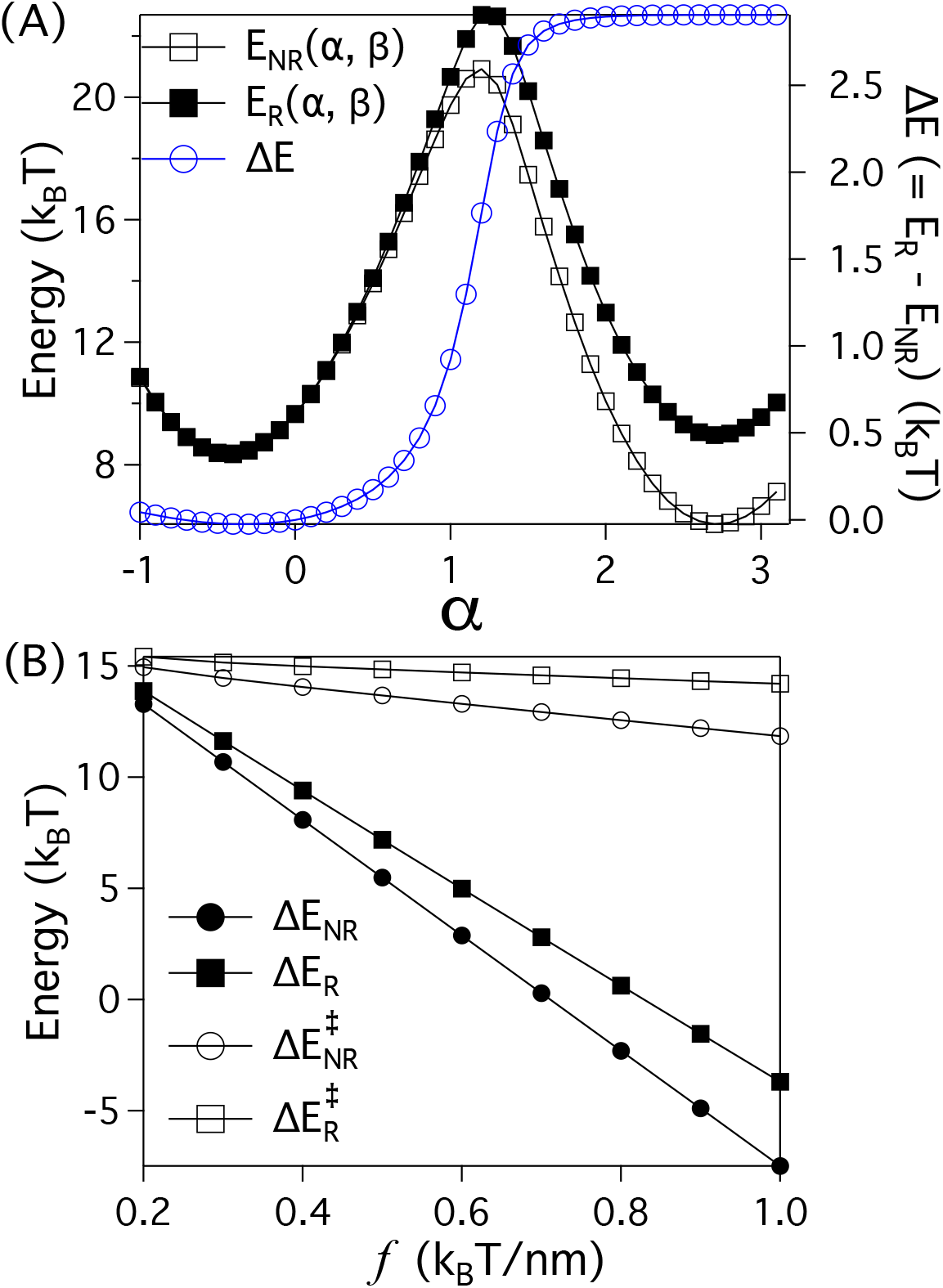
(A) Plot of *E_R_*(*α*, *β*) (solid squares), *E_NR_*(*α*, *β*) (empty squares) and Δ*E*(= *E_R_*(*α*, *β*) – *E_NR_*(*α*, *β*)) (blue empty circles) as a function of *α*. Due to the HPC rotation, the barrier for DNA unwrapping increases by ≈ 2.5 *k*_B_*T*. (B) Δ*E_R_*(= *E_R_*(*α* ≈ *π*) – *E_R_*(*α* ≈ 0)) (solid squares) and Δ*E_NR_*(= *E_NR_*(*α* ≈ *π*) – *E_NR_* (*α* ≈ 0)) (solid circles) as a function of *f*. 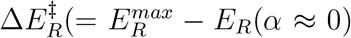) (empty squares) and 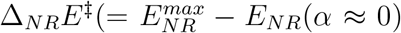) (empty circles) as a function of *f*. Here, 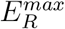 and 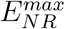 is the maximum (corresponds to the transition region) in the energy during the transition from *α* ≈ 0 to *α* ≈ *π* state. Note that if the dimensions of the cylinder and the value of *α* are known, then the HPC rotation angle can be calculated using Eq. (S8) in the SI. The HPC parameters are *R* = 4.2 nm, *H* = 2.5 nm, *l_p_* = 50 nm, *ϵ_ads_* = 0.7 *k*_B_*T*, *f* = 0.8 *k*_B_*T*/nm.

It is important to note that the calculated increase in the barrier using Eq. 2 is a lower bound. In addition to the purely energetic barrier arising from the elastic model there is a penalty (*E_dyn_* ∝ *ηRH*, where *η* is the viscosity of water) to rotate the cylinder through the viscous medium. The resulting kinetic barrier would further enhance the total barrier that must be surmounted to unwrap the outer turn.

## Discussion

We used constant force simulations using the coarse-grained SOP model, without any adjustable parameter, to determine the structural transitions that take place during unwrapping of human *α*-satellite DNA sequence from the nucleosome as well as in the assembly process upon quenching the force. Our simulations, using a single bead resolution of the 1,268 combined number of nucleotides and residues of histone proteins, is consistent with the two stage unwrapping transition observed in several experiments. The length of DNA released in the two transitions are in excellent agreement with single molecule pulling experiments. ^12^ The global two stage transition is found to be nearly independent of DNA sequence (experiments and simulations often use the Widom high affinity nucleosome 601 sequence), implying that certain aspects of unwrapping depends only on the nucleosome structure. This is similar to the finding that native topology determines the force induced unfolding pathways of globular proteins.^43^

### Energetics and time scales of outer turn unwrapping

The critical force, *f_cI_*, for the first reversible transition (centered at approximately at ≈ 3 pN or ≤ 6 pN in experiments^12,14^ and ≈ 5 pN in our simulations) is needed to overcome the interactions of the outer turn of the DNA with histones. The lack of dependence of the outer turn rip on the pulling direction (Figure 4) confirms that outer loop of the DNA reversibly wraps and unwraps around the HPC. We conclude that the apparent barrier (≈ 15 *k*_B_*T*), not that dissimilar to the energy required to unbind the outer turn, ^22^ which we take to be the minimum free energy barrier. Our estimate for the barrier height when combined with the attempt frequency^30^ of *k*_0_ ≈ 4.0 × 10^6^ yields a rupture time for outer turn (as per Kramers theory) on the order of 800 ms at zero force, which is not inconsistent with the results in Ref.^14,44,45^

We note that the estimates of outer turn unwrapping time (lifetime of the fully wrapped state) seems to vary widely, ranging from about 250 ms using FRET and time-resolved SAXS experiments^44,45^ to ten seconds or more using pulling experiments (see Fig. 4b in Ref.^13^ for comparison between three data sets). Although experiments using mechanical forces have provided considerable insights into the disassembly of single nucleosomes, it is unclear if extrapolation using *f* ≠ 0 measurements to *f* = 0 to extract the lifetimes of the wrapped or unwrapped states is reasonable. More importantly, the validity of using extension as the sole reaction coordinate for a complex DNA-protein complex is unclear. Even for the simpler cases of force-induced unfolding of globular proteins and RNA, extension is often inappropriate.^46^ Nevertheless, considering the uncertainties in the different experimental techniques and protocols^13,44,45^ and the limitations of theory and simulations, it is likely that the time scale for outer unwrapping transition, at zero force, is likely to be between 250 ms to about a second.

### Rupture of the inner turn

The second transition in which the inner loop of the DNA unwraps is to a large extent cooperative (see Figure 4), which accords well with pulling experiments using magnetic tweezers.^47^ This transition, which could have short-lived intermediates at high pulling forces (see *R_ee_*(*t*) plot in Figure 2A), is not reversible,^12^ thus making it difficult to estimate the stability and free energy barrier to rupture. The mechanical energy needed for this transition is ≈ 75 *k*_B_*T*, which is substantial. The minimum estimate for this transition from experiments is ≈ 45 *k*_B_*T*, which is the product of the unbinding force (≈ 9 pN) and length gain (≈ 21 nm). Our simulations show that for the inner turn to unwrap, sufficient torque to rotate the HPC by 180°, is required in order to overcome the DNA-histone interactions. The presence of histone tails on either side of the DNA prevent the rotation of the HPC by 180° (Figure 1D), which should increase the mechanical energy needed to disrupt the DNA-HPC interaction.

The rotational barrier has two contributions. One is energetic, which we could using the KS elastic model (Figure 7). The other is kinematic in origin that arises because of the rotation of the HPC moment of inertia. The kinetic barrier is ∝ *ηω* where *ω* is the angular velocity. The kinematic consequences affect wrapping and unwrapping in totally different ways, leading the asymmetry in the role of histone rotation in the rupture process.

### Forced-unwrapping and force-quench assembly

Our simulations show that there is a fundamental difference in unwrapping and reassembly upon force quench. In forced-unwrapping, the torque needed to rotate the HPC comes from force, which facilitates the HPC rotation in peeling the inner turn. When force is quenched (reduced in the release cycle in optical tweezer experiments), the principal moment of inertia of the HPC has to align along the force direction in order for DNA to wrap around the HPC. In contrast to the “directed” rotation in the unwrapping process, the rotation in the release cycle occurs stochastically. In other words, the external force does not play any role in aligning the HPC. Simulations show that the folding and the unfolding pathways of the DNA from the HPC during the second transition are different. The difference in the role played by the external force in DNA unwrapping and rewrapping explains the hysteresis observed in experiments.

### Prospects for experiments

The major findings of this work are that the histone core rotates during the unwrapping of the second turn in the presence of force as well during the assembly upon force quench from the fully unwrapped state. It is worth restating that HPC rotation plays an entirely different role in the unwrapping and rewrapping processes. When the inner turn is mechanically forced to unwrap, a large force is needed to rotate the HPC in almost deterministic manner (fluctuations in *R_ee_*(*t*) are suppressed). Two experiments might shed light on our predictions. (1) To observe histone rotation it will be necessary to perform three color FRET experiments by labeling two residues on the protein and one on a base pair in the inner turn. Combined force and multiply labelled FRET could be used to discern HPC rotation. We are unsure if such experiments are feasible. (2) The theoretical model suggests that that the time for unwrapping of the inner wrap would increase as exp(*cη*), where c is a constant. Thus, small changes in *η* are predicted to have a large effect on the rate of unwrapping. Naively, it appears that the frictional effect of rotation may be easier to test in experiments.

## Conclusion

Our simulations show that forced unwrapping and force quench rewrapping of DNA is coupled to histone rotation, which has not been sufficiently emphasized. It is unclear if the response of nucleosomes to force would differ if lysine at various locations (K9 or K27) in the histones are modified. The SOP model can be used to probe the consequences of such post-translational modifications.

We will be remiss if we did not point out the limitations of the study. (i) We have already stated that the duration of the simulations is much shorter than in experiments. Nevertheless, it is surprising that both the length gains in DNA upon unwrapping and the mean forces for rupture are in reasonable agreement with experiments. (ii) We have not explicitly considered hydrogen bond interactions between the nucleotides because the model ignores the DNA sequence, which plays an important role. It will be important to investigate consequences of the sequence-dependent elasticity of DNA^48^ although global features of unwrapping (two stage unwrapping and HPC rotation) are likely to be robust.

## Acknowledgements

We are grateful to Debayan Chakraborty, Sucheol Shin, and Guang Shi, and Helmut Schiessel for useful discussions. Much of this work was done while the authors were in the Institute for Physical Science and Technology in the University of Maryland. This work was supported by NIH (GM - 107703), and the Welch Foundation Grant F-0019 through the Collie-Welch chair. G.R. acknowledges support from the National Supercomputing Mission (NSM) through the grant MeitY/R&D/HPC/2(1)/2014.

