## Supplementary Information for "Asymmetry in Histone Rotation in Forced Unwrapping and Force Quench Rewrapping in a Nucleosome"

### Supporting Information for "Asymmetry in Histone Rotation in Forced Unwrapping and Force Quench Rewrapping in a Nucleosome"

Govardhan Reddy[†] and D. Thirumalai[*‡]

†*Solid State and Structural Chemistry Unit,*

*Indian Institute of Science, Bangalore, Karnataka, India 560012*

[‡]*Department of Chemistry, The University of Texas, Austin, TX 78712*

(Dated: October 20, 2020)

### I. METHODS

**Self Organized Polymer (SOP) Model for the Nucleosome:** The SOP model[1] for the nucleosome is constructed using the crystal structure[2] of the human nucleosome available in the Protein Data Bank (PDB ID: 2CV5). The 146 base pair corresponds to the palindromic $\alpha$-satellite DNA sequence. Residues in the histone protein and the nucleotides in the DNA are represented using a single bead. The total energy of the nucleosome is a sum of bonded (B) and non-bonded (NB) interactions, where the NB interactions are a sum of native (N) and non-native (NN) interactions. Interaction between two beads (separated by at least two other beads along the sequence if they are on the same chain) is deemed to be native if the distance between them is less than a cutoff value, $R_c$, in the coarse grained representation of the crystal structure. Each strand in the DNA has 147 base pairs (bps), and the number of residues in the histone proteins H3, H4, H2A and H2B with the disordered tails is 136, 103, 130 and 126, respectively. The residues in the histone tails are 1-44 in H3, 1-30 in H4, 1-26 in H2A, and 1-34 in H2B. In total, there are 1,268 (1,000) beads in the nucleosome with (without) histone tails. Because this is a large complex it is necessary to use coarse grained models in simulating the effect of force on a nucleosome.

The disordered tails that are not resolved in the crystal structure are primarily located in the N-terminal regions of H3, H4, H2A and H2B. To understand the role of the tails in the DNA unwrapping from the Histone Protein Core (HPC), we performed simulations with and without histone tails. The SOP energy function for the nucleosome without the histone tails is,

$$
\begin{aligned}
E_T = & -\sum_{X=1}^{N^{HPC}+N^{DNA}} \left( \sum_{i=1}^{N_B^X} \frac{k}{2} R_o^2 \log\left(1 - \frac{(r_i^X - r_{cry,i}^X)^2}{R_o^2}\right) + \sum_{i=1}^{N^X-2} \epsilon_l \left(\frac{\sigma_{i,i+2}^X}{r_{i,i+2}^X}\right)^6 \right) \\
& + \sum_{Y=HPC,DNA,HPC-DNA} \left( \sum_{i=1}^{N_N^Y} \epsilon_h^Y \left[ \left(\frac{r_{cry,i}^Y}{r_i^Y}\right)^{12} - 2\left(\frac{r_{cry,i}^Y}{r_i^Y}\right)^6 \right] + \sum_{i=1}^{N_{NN}^Y} \epsilon_l \left(\frac{\sigma_i^Y}{r_i^Y}\right)^6 \right),
\end{aligned} \tag{S1}
$$

where $N^{HPC}(=8)$ and $N^{DNA}(=2)$ are the number of histone chains in the HPC and DNA chains in the nucleosome, respectively. In the chain $X$, which can be either a protein or a DNA, $N^X$ is the number of beads in the chain, $N_B^X$ is the number of bonds, $r_i^X$ is the distance between the $i^{th}$ pair of beads that have a bond between them, $r_{cry,i}^X$ is the corresponding distance in the crystal structure. In the above equation, $\sigma_{i,i+2}^X$ is the sum of the radii of the beads $i$ and $i+2$, and $r_{i,i+2}^X$ is the distance between the beads $i$ and $i+2$. In Eq. S1, $Y$

represents the interaction between the pairs of beads that belong to the HPC or pairs of beads that belong to the DNA or a bead from the HPC and the other bead from the DNA. In Eq. S1, $N_N^Y$ and $N_{NN}^Y$ represent the total number of native and non-native interactions, respectively, $r_i^Y$ is the distance between the $i^{th}$ pair of beads that interact through native or non-native potentials, and $r_{cry,i}^Y$ is the corresponding distance in the crystal structure, $\sigma_i^Y$ is the sum of the radii of the $i^{th}$ pair of beads. The values of the interaction parameters in the energy function are given in Table S1. The reasonable agreement demonstrated here and the predictions made previously[3] shows that the SOP model for protein-DNA interactions is transferable.

In order to assess the importance of the histone tails, we also performed simulations using the SOP model for the full structure of the nucleosome. The histone tails interact with the rest of the histone core and the DNA via excluded volume and electrostatic interactions. We model the interactions between the charged residues (Arg, Asp, Glu, His and Lys) with the DNA using a screened Coulomb potential[3,4]. The total energy of the nucleosome with the histone tails is,

$$E_T^{Tail} = E_T + \sum_{i=(Arg,Asp,Glu,His,Lys)} \sum_{j=1}^{N^{DNA} \times N_L^{DNA}} \frac{q_i q_j}{4\pi\epsilon r_{ij}} e^{-\kappa r_{ij}}, \tag{S2}$$

where, $N_L^{DNA}(= 147)$ is the length of the DNA, $q_i$ and $q_j$ are the charges on the charged residues in the histone tail and DNA, respectively. The charges are at the center of the respective beads. The charge on the DNA is negative, $q_j = -|e|$, where $e$ is the charge of an electron. The value of the dielectric constant, $\epsilon$, in the simulations is $10\epsilon_o$ ($\epsilon_o$ is vacuum permittivity), and $\kappa$, the inverse Debye length, is calculated for a monovalent 100 mm salt concentration.

**Simulations:** In order to dissect the kinetics of force-induced unwrapping of nucleosome and reassembly (upon force quench) we used Brownian dynamics simulations. The equations of motion were integrated using the Ermak and McCammon algorithm[5] without hydrodynamic interactions. In our simulations, length, mass and friction coefficient are fixed. Without loss of generality we set length, $a = 1$ Å , energy, $\epsilon_l = 1$ kcal/mol, and the friction coefficient $\zeta = 1.8 \times 10^{-10}$ g/s. The friction coefficient for the bead in the protein, $\zeta^P = 50\zeta$, and for the DNA bead, $\zeta^{DNA} = 100\zeta$. Because of the size difference between the nucleotide and protein residue, $\zeta^{DNA} > \zeta^P$. The characteristic time scale in the simulation, $\tau_L$ is estimated using $a$, $\epsilon$ and $\zeta$. The friction coefficient for a protein bead in water[6] $\zeta^{prot} \approx 9 \times 10^{-9}$ g/s, which implies $\zeta = 1.8 \times 10^{-10}$ g/s. The time scale $\tau_L = \zeta a^2/\epsilon = 0.26$ ps. Simulations are performed at temperature, $T = 300$ K. The equations of motion are integrated using a time step of $\Delta t = 0.1\tau_L$. In the constant force simulations, one end of the double stranded DNA, which wraps the HPC is fixed and a constant $f$ is applied to the other end (Figure S1).

**Quantifying HPC rotation:** The rotation of HPC due to the torque generated by the application of mechanical force, $f$, is quantified using the angle, $\theta$, between the end-to-end vector of the DNA, $\vec{R_{ee}}$, and the principal moment of inertia of HPC, $\vec{I}$, which roughly points along the nucleosome super helical axis. The angle $\theta$ is defined as,

$$\cos(\theta) = \frac{\vec{R_{ee}} \cdot \vec{I}}{|\vec{R_{ee}}||\vec{I}|}. \tag{S3}$$

$\vec{R_{ee}}$ of the dsDNA is given by $\vec{R_{ee}} = (\vec{R^1_{ee}} + \vec{R^2_{ee}})/2$, where $\vec{R^1_{ee}}$ and $\vec{R^2_{ee}}$ are the end-to-end vectors of the 2 DNA strands in the dsDNA. The eigenvector, $\vec{I}$, corresponds to the maximum eigenvalue of the moment of inertia tensor of the HPC, $S^{HPC}$. The $xy$ component of the moment of inertia tensor of the HPC, $S^{HPC}_{xy}$, in Cartesian coordinates is given by,

$$S^{HPC}_{xy} = \frac{1}{2N^2_{HPC}} \sum_{i=1}^{N_{HPC}} \sum_{j=1}^{N_{HPC}} (x_i - x_j)(y_i - y_j), \tag{S4}$$

where $N_{HPC} = 974$ (706) is the number of coarse-grained protein beads in the HPC with (without) the histone tails, $x_i$ and $y_i$ are the $x$ and $y$ components of the position vector of protein bead $i$ in the Cartesian coordinates. In computing $S^{HPC}$, the mass of all the coarse-grained protein beads are taken to be unity. To quantify the rotation using $\theta$ (Eq. S3), we computed $\vec{I}$ as a function of time in the DNA unwrapping simulation trajectory using the program visual molecular dynamics (VMD)[7].

**Theoretical Model:** In order to calculate the increase in the barrier due to rotation of the HPC, which is especially dominant in the second stage of the unwrapping transition (see the main text for details), we use the Kulic-Schiessel (KS) model[8] based on DNA elasticity. The bending elastic energy of the WLC is given by,

$$E_{bend} = \frac{A}{2} \int_0^L \kappa^2(s) \, \mathrm{d}s, \tag{S5}$$

where $A = l_p k_B T$. In Eq. S5, $l_p$, $k_B$ and $T$ are the persistence length, Boltzmann constant, and temperature, respectively. Following the notation in KS, $f$ is applied to the WLC along the $y$-axis (Figure S1). As a result, the WLC bends and desorbs from the cylinder and simultaneously the cylinder, representing the HPC, rotates to align along the force axis, as explicitly illustrated in our simulations (see the main text). The desorption angle $\alpha$ describes the amount of the WLC adsorbed onto the cylinder ($\alpha = 0$, and $\pi$ corresponds to one turn of wrapped DNA (1.0**N**), and WLC fully unwrapped DNA, respectively). The angle $\beta$ describes the degree of rotation of the cylinder representing the HPC, and is the angle between the axis of the cylinder, $\hat{n}$, and the $z$-axis (Figure S1). Due to the symmetry, the cylinder, $\hat{n}$, rotates only in the $yz$ plane.

The total energy of the nucleosome in the KS model is,

$$E_R(\alpha, \beta) = E_{NR}(\alpha, \beta) + E_{rot} = 2(R\alpha\epsilon_{ads} + E_{bend} - f\Delta y) + E_{rot}, \tag{S6}$$

where $E_{NR}(\alpha, \beta) = 2(R\alpha\epsilon_{ads} + E_{bend} - f\Delta y)$. The factor of 2 in Eq. S6 is a consequence of symmetry. The first term represents the adsorption energy, the second term describes the bending penalty, the third term is the mechanical energy due to the stretching force, and the fourth term ($E_{rot}$) describes the energy required to rotate the solid cylinder (HPC) by the torque arising from $f$.

KS estimated[8,9] $E_{bend}$ and $\Delta y$ using three approximations: (i) entropic shape fluctuations of the DNA are neglected, which is valid at large $f$ or low $T$, (ii) DNA arms are asymptotically straight and parallel to the $y$-axis (the force axis), (iii) the length of the DNA arm is big, $L/\lambda \gg 1$, where $\lambda = \sqrt{\frac{l_p k_B T}{f}}$. All of these approximations are reasonable but require scrutiny, which our simulations provide. With these approximations, the total Hamiltonian can be written[8,9] as,

$$\begin{aligned} E_R(\alpha, \beta) &= E_{NR}(\alpha, \beta) + E_{rot} \\ &= 2R\alpha\epsilon_{ads} + 2fR\left(\cos\beta\sin\alpha - \frac{H}{2\pi R}(\pi - \alpha)\sin\beta - \alpha\right) \\ &\quad + 8\sqrt{l_p k_B T f}\left(1 - \sqrt{\left(1 + \frac{R}{\overline{R}}\cos\beta\cos\alpha + \frac{H}{2\pi\overline{R}}\sin\beta\right)/2}\right) + E_{rot}. \end{aligned} \tag{S7}$$

We discovered in our simulations that HPC rotation plays an important role, especially in the second stage of nucleosome unwrapping, which would increase the energetic barrier by $E_{rot}$. The constraint that the torque at the ends of the DNA arms should vanish gives a relation between $\alpha$, $\beta$ and $f$,

$$\frac{\sqrt{2}\lambda}{R}\sqrt{\frac{1-\left(\frac{R}{\overline{R}}\cos\beta\cos\alpha+\frac{H}{2\pi\overline{R}}\sin\beta\right)}{\sin^2\alpha+(\sin\beta\cos\alpha-\frac{H}{2\pi R}\cos\beta)^2}}\left(\sin\beta\cos\alpha-\frac{H}{2\pi R}\cos\beta\right)$$
$$=\sin\beta\sin\alpha+\frac{H}{2\pi R}(\pi-\alpha)\cos\beta. \tag{S8}$$

We estimated the contribution of the barrier to rotation, $E_{rot}$, purely from the elastic model, which increases the transition energy barrier for the nucleosome to go from the state where one turn of DNA is wrapped on the HPC ($\alpha = 0$) to the state where DNA is completely unwrapped ($\alpha = \pi$). The energy required to rotate the cylinder through an angle $\beta$ by the torque, $\tau(\alpha, \beta')$, generated by $f$ acting on the WLC arms subject to the constraint (Eq. S8) relates $f$, $\alpha$, and $\beta'$. We find that $E_{rot}$ is given by,

$$E_{rot}(\alpha,\beta) = -\int_0^{\beta}\tau(\alpha,\beta')\ \mathrm{d}\beta'. \tag{S9}$$

To estimate $\tau(\alpha, \beta)$ in the cylinder orientation specified by $\alpha$ and $\beta$, it is necessary to describe the path of the WLC on the cylinder. Since the WLC takes a helical path on the cylinder, for angles, $\alpha$ and $\beta$=0, the path of the WLC is given by,

$$\vec{h}(t) = \begin{pmatrix} R\cos t \\ R\sin t \\ \frac{H}{2\pi}(\pi - t)\end{pmatrix} \tag{S10}$$

for $\alpha < t < (2\pi - \alpha)$. The path of the WLC on the cylinder with non-zero orientation $\beta$ is obtained by multiplying $\vec{h}(t)$ with the rotation matrix to give

$$\vec{h}(t,\beta) = \begin{pmatrix} 1 & 0 & 0 \\ 0 & \cos\beta & -\sin\beta \\ 0 & \sin\beta & \cos\beta \end{pmatrix} \times \begin{pmatrix} R\cos t \\ R\sin t \\ \frac{H}{2\pi}(\pi - t)\end{pmatrix} = \begin{pmatrix} R\cos t \\ R\cos\beta\sin t - \frac{H}{2\pi}(\pi-t)\sin\beta \\ R\sin\beta\sin t + \frac{H}{2\pi}(\pi-t)\cos\beta \end{pmatrix}. \tag{S11}$$

The unit tangent vector, $\hat{h}'(t, \beta)$ is given by,

$$\frac{1}{\overline{R}}\begin{pmatrix} -R\sin t \\ R\cos\beta\cos t + \frac{H}{2\pi}\sin\beta \\ R\sin\beta\cos t - \frac{H}{2\pi}\cos\beta \end{pmatrix} \tag{S12}$$

where $\overline{R} = \sqrt{R^2 + (\frac{H}{2\pi})^2}$. The WLC arms leave the cylinder at positions $\vec{h}(\alpha, \beta)$ and $\vec{h}(2\pi - \alpha, \beta)$, and the force $f$ acts on the cylinder at these positions along the unit tangent vectors

$\hat{h}'(\alpha, \beta)$ and $\hat{h}'(2\pi - \alpha, \beta)$. Thus, the expression for the torque is

$$\begin{aligned}\tau(\alpha,\beta) &= (\vec{h}(\alpha,\beta) \times f\hat{h}'(\alpha,\beta)) - (\vec{h}(2\pi-\alpha,\beta) \times f\hat{h}'(2\pi-\alpha,\beta)) \\ &= -\frac{fHR}{\pi \overline{R}}[\sin\alpha + (\pi - \alpha)\cos\alpha]\ \hat{x}.\end{aligned} \tag{S13}$$

We can calculate $E_{rot}(\alpha, \beta)$ using Eq. S8, S9, S13 and rewriting Eq. S9 as,

$$E_{rot}(\alpha,\beta) = -\int_0^{\alpha} \tau(\alpha',\beta)\left(\frac{\mathrm{d}\beta}{\mathrm{d}\alpha'}\right)\mathrm{d}\alpha'. \tag{S14}$$

It is possible to evaluate Eq. S14 numerically. The enhanced rotational barrier, given in Eq. S14 arises solely from energetic considerations. However, the cylinder, representing the HPS, rotates in water, which yields an additional purely kinetic barrier. The value of the latter is determined by the solvent viscosity, and other characteristics of the cylinder. Thus, $E_{rot}$ is a lower bound to the barrier due to rotation of the HPC in water.

---

[9] Igor M. Kulic, Herve Mohrbach, Rochish Thaokar, and Helmut Schiessel. Equation of state of looped DNA. *Phys. Rev. E*, 75, 2007.

**FIGURES**

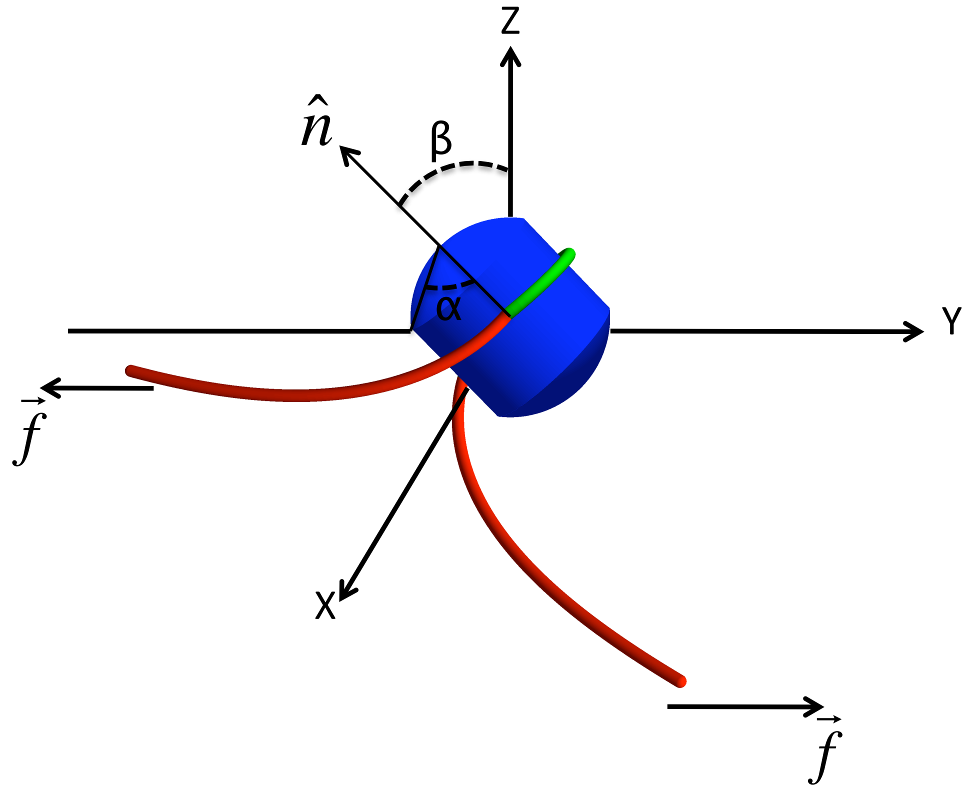


FIG. S1. Schematic of the nucleosome under tension[8]. The HPC is depicted as a cylinder in blue. The DNA adsorbed on the HPC is in green and the DNA arms are in red. Angle $\alpha$ is the desorption angle of the DNA and angle $\beta$ is the degree of rotation of the HPC and is given by the angle between the HPC axis, $\hat{n}$, and the $z$-axis.

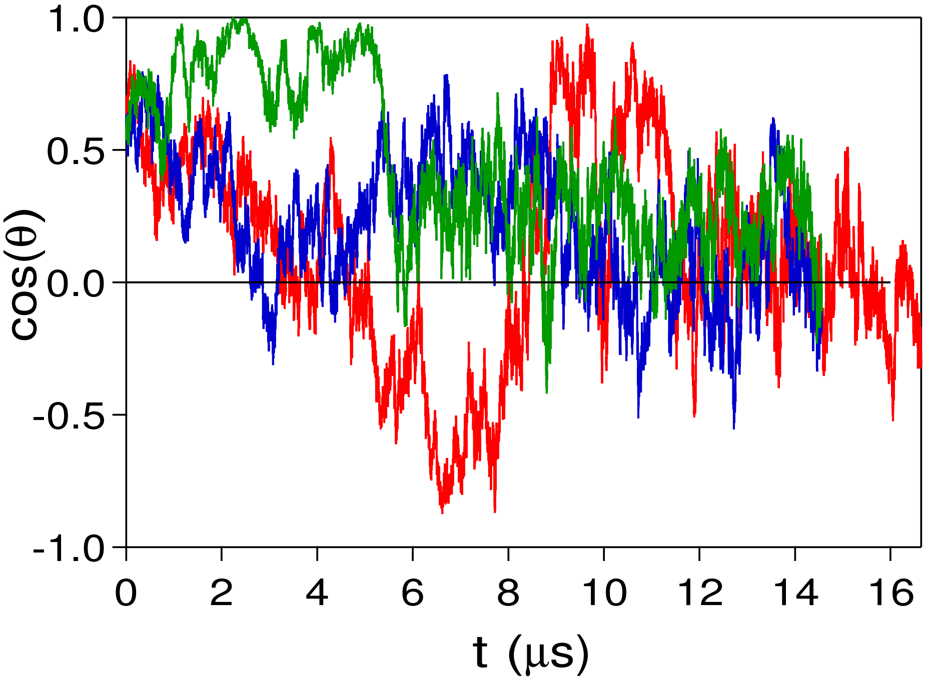


FIG. S2. Plot of $\cos(\theta)$ (Eq. S3) as a function of $t$ for three trajectories (red, green, and blue) recorded during rewrapping after a force quench to $f_Q = 0$. The HPC orientation fluctuates randomly as the DNA wraps around the HPC in contrast to the unwrapping pathway where the HPC rotates by 180° (see the results in the main text).

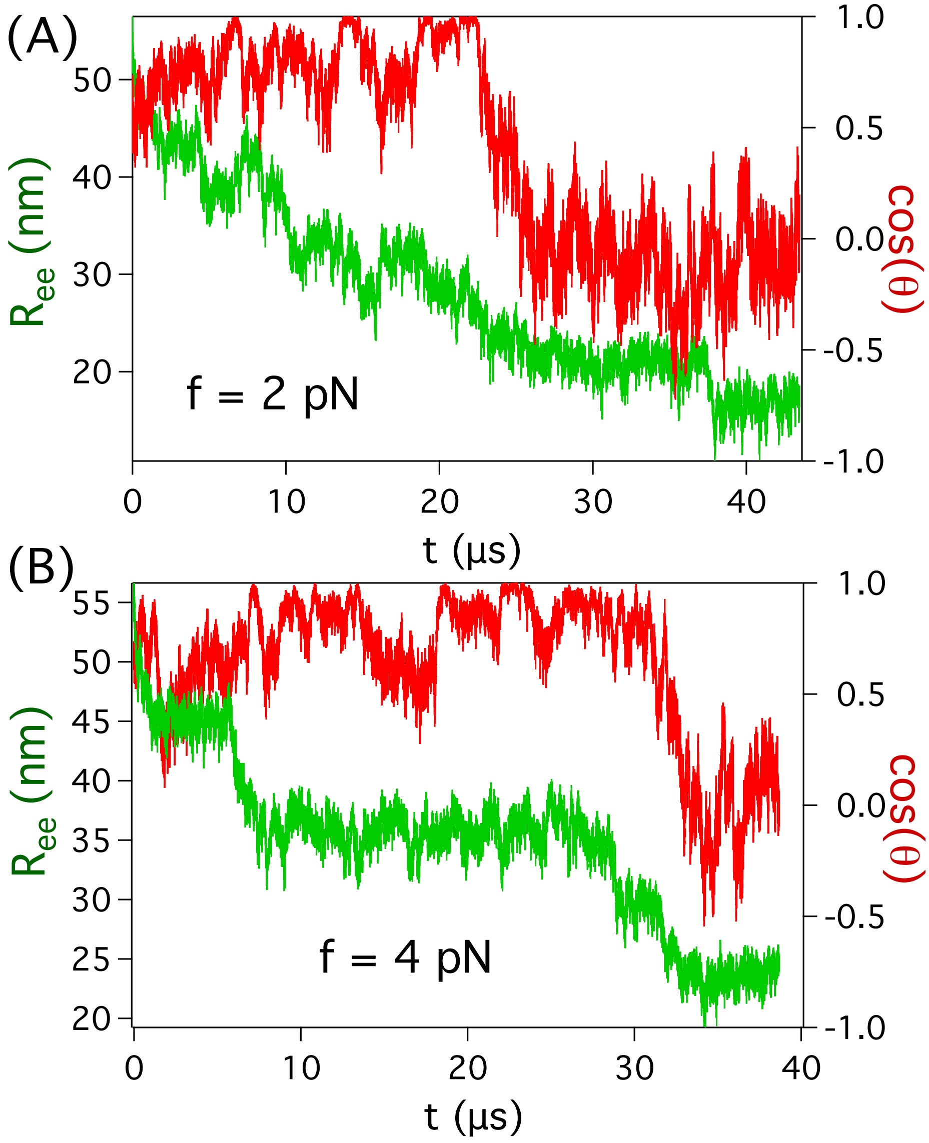


FIG. S3. Rewrapping dynamics. $R_{ee}$ and $\cos(\theta)$ are plotted as a function of $t$ for a DNA folding trajectory for (A) $f_Q = 2$ pN, and (B) $f_Q = 4$ pN.

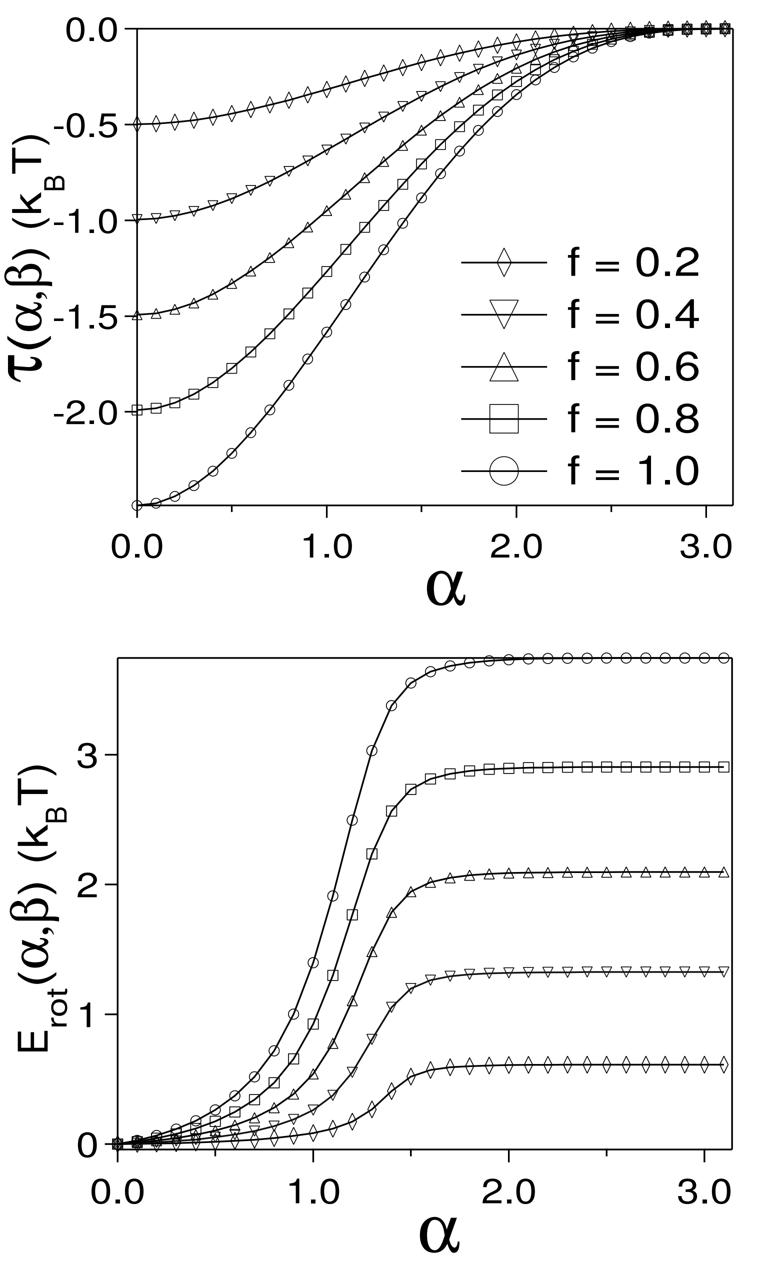


FIG. S4. (A) Plot of $\tau(\alpha, \beta)$ as a function of $\alpha$ for different force, $f$, values ($f$ is in units of $k_{\rm B}T/nm$) subject to the constraint between $\alpha$, $\beta$ and $f$ (Eq. S8). (B) $E_{rot}(\alpha, \beta)$ as a function of $\alpha$ for different $f$. Similar symbols in (A) and (B) represent identical forces. The other parameters are $R = 4.2$ nm, $H = 2.5$ nm, $l_p = 50$ nm, and $\epsilon_{ads} = 0.7\ k_{\rm B}T$.

**TABLES**

TABLE S1. Parameters for the SOP model of mononucleosome without the histone tails (model-1).

| | DNA | HPC | HPC-DNA |
|---|---|---|---|
| $R_o$ | 2.0 | 2.0 | - |
| $k$ | 20 kcal/(mol. Å$^2$) | 20 kcal/(mol. Å$^2$) | - |
| $R_c$ | 14 Å | 8 Å | 11 Å |
| $\epsilon_h^Y$ | 0.7 kcal/mol | 2.0 kcal/mol | 1.18 kcal/mol |
| $\epsilon_l$ | 1.0 kcal/mol | 1.0 kcal/mol | 1.0 kcal/mol |
| $\sigma^Y$ | 7.0 Å | 3.8 Å | 5.4 Å |
| $\sigma^X$ | 7.0 Å | 3.8 Å | - |
| $\zeta$ | 100 $\tau_L^{-1}$ | 50 $\tau_L^{-1}$ | - |
| $N_N^Y$ | 1306 | 2123 | 266 |
| $N_{NN}^Y$ | 41183 | 245354 | 207298 |
